# Allosteric inhibition of CXCR1 and CXCR2 abrogates Th2/Th17-associated Allergic Lung Inflammation in Mice

**DOI:** 10.1101/2024.05.13.593638

**Authors:** Koa Hosoki, Annamalai Govindhan, John M Knight, Sanjiv Sur

**Author notes:** Corresponding author: Sanjiv Sur, Address: Baylor College of Medicine, 1 Baylor Plaza, Houston, TX 77030.

## Abstract

**Background:** IL4, IL5, IL13, and IL17-producing CD4 T helper 2 (Th2)-cells and IL17-producing CD4 T helper 17 (Th17)-cells contribute to chronic eosinophilic and neutrophilic airway inflammation in asthma and allergic airway inflammation. Chemokines and their receptors are upregulated in Th2/Th17-mediated inflammation. However, the ability of CXCR1 and CXCR2 modulate Th2 and Th17-cell-mediated allergic lung inflammation has not been reported.

**Methods:** Mice sensitized and challenged with cat dander extract (CDE) mount a vigorous Th2-Th17-mediated allergic lung inflammation. Allosteric inhibitor of CXCR1 and CXCR2, ladarixin was orally administered in this model. The ability of ladarixin to modulate allergen-challenge induced recruitment of CXCR1 and CXCR2-expressing Th2 and Th17-cells and allergic lung inflammation were examined.

**Results:** Allergen challenge in sensitized mice increased mRNA expression levels of *Il4, Il5, Il13, Il6, Il1β, Tgfβ1, Il17, Il23, Gata3,* and *Rorc*, and induced allergic lung inflammation characterized by recruitment of CXCR1- and CXCR2-expressing Th2-cells, Th17-cells, neutrophils, and eosinophils. Allosteric inhibition of CXCR1 and CXCR2 vigorously blocked each of these pro-inflammatory effects of allergen challenge. CXCL chemokines induced a CXCR1 and CXCR2-dependent proliferation of IL4, IL5, IL13, and IL17 expressing T-cells.

**Conclusion:** Allosteric inhibition of CXCR1 and CXCR2 abrogates blocks recruitment of CXCR1- and CXCR2-expressing Th2-cells, Th17-cells, neutrophils, and eosinophils in this mouse model of allergic lung inflammation. We suggest that the ability of allosteric inhibition of CXCR1 and CXCR2 to abrogate Th2 and Th17-mediated allergic inflammation should be investigated in humans.

## Introduction

Asthma is a chronic inflammatory disease of the airways characterized by chronic eosinophilic and neutrophilic allergic inflammation. IL4-, IL5-, and IL13-producing CD4 T helper 2 (Th2)-cells^1^ and IL17-producing Th17-cells^2^ contribute to this inflammation. IL4 stimulates Th2 differentiation of CD4+ T-cells by upregulating its master regulator GATA3^3^, and these Th2-cells induce allergic lung inflammation^3,4^. IL23, IL1β, and IL6 stimulate CD4+ T-cells to differentiate toward Th17^5–7^ by upregulating its master regulator retinoic acid–related orphan receptor-gamma t (RORγt)^8^, Th17-cells mediate neutrophilic inflammation^9^.

A large volume of literature indicates that CC-chemokine receptors (CCR) are present on eosinophils, Th2-cells, and Th17-cells, and modulate eosinophilic and neutrophilic airway inflammation in asthma by attenuating chemokine-induced migration and activation Th2-associated inflammatory immune cells^10^. CCR3 is expressed on eosinophils and regulates recruitment and degranulation of eosinophils^11^. CCR3, CCR4, and CCR8 are expressed on Th2-cells^12^, and CCR4 plays a role in recruiting Th2-cells^13,14^. CCR3 expressing eosinophils, and CCR4 and CCR8 expressing Th2-cells can be detected in endobronchial biopsies performed in asthmatic subjects after allergen challenge^15^. Th17-cells express CCR4, CCR5, and CCR6^16–18^, and CCR6 facilitates recruitment of Th17-cells^18^.

Unlike the plethora of studies linking CCR receptors with allergic inflammation, less is known about the role of CXC-chemokine receptors (CXCR) in allergic inflammation. This is somewhat surprising because the levels of CXCL chemokines are elevated in the airways in asthma^19–21^, and the expression of CXCL1, CXCL5, and CXCL8 in bronchoalveolar lavage fluid (BALF) and endobronchial biopsies is higher in subjects with asthma than healthy control subjects^20,21^. Furthermore, the levels of CXCL5 and CXCL8 are higher in patients with acute severe asthma^19,20^. CXCR1 and CXCR2 are G protein-coupled chemokine receptors for CXCL1, CXCL2, CXCL3, CXCL5, CXCL6, CXCL7, and CXCL8, and are expressed on many immune and lung structural cells^22–24^. We reported that administration of CXCR2 small molecule inhibitor SB225002 and dual CXCR1 and CXCR2 inhibitor reparixin suppressed allergic airway inflammation and serum IgE levels^25,26^. However, the role of CXCR1 and CXCR2 in modulating Th2 and Th17-associated allergic lung inflammation has not been reported. In the present study, we examined the role of allosteric inhibition of CXCR1 and CXCR2 in Th2 and Th17-associated allergic lung inflammation.

## Materials and methods

### Allergenic extracts

Lyophilized cat dander extract (CDE) (lots#253320, 351876, and 392583) was purchased from Greer Labs (Lenoir, NC). The level of endotoxin in CDE was measured using a LAL chromogenic endotoxin quantitation kit (Thermo Scientific, Hudson, NH), and was less than 0.1 pg/µg CDE protein, hence unlikely to contribute significantly to inflammation^27^.

### Protocols used for animal studies

C57Bl/6 mice were anesthetized with an intraperitoneal injection of a low dose of xylazine/ketamine anesthetic mixture for intranasal administration of CDE and sacrificed by a lethal dose of intraperitoneal xylazine/ketamine. The protocol was approved by the IACUC of Baylor College of Medicine.

### CDE Multiple Challenge Model (CDE-MCM) to induce allergic sensitization

Naïve wild type (WT) mice were sensitized by five intranasal challenges of CDE (100 µg/60µl) on days 0, 1, 2, 3, and 4. After a rest period of 7 days, these mice were challenged with an intranasal dose of CDE or phosphate-buffered saline (PBS) on day 11 and sacrificed at 2 h, 4 h,16 h, 28 h, 40 h, and 72 h post-CDE challenge. Some mice challenged with CDE on day 11 were orally treated with 15 mg/kg body weight of ladarixin on days 11, 12, and 13 (**Figure 1A**).

**Figure 1.**
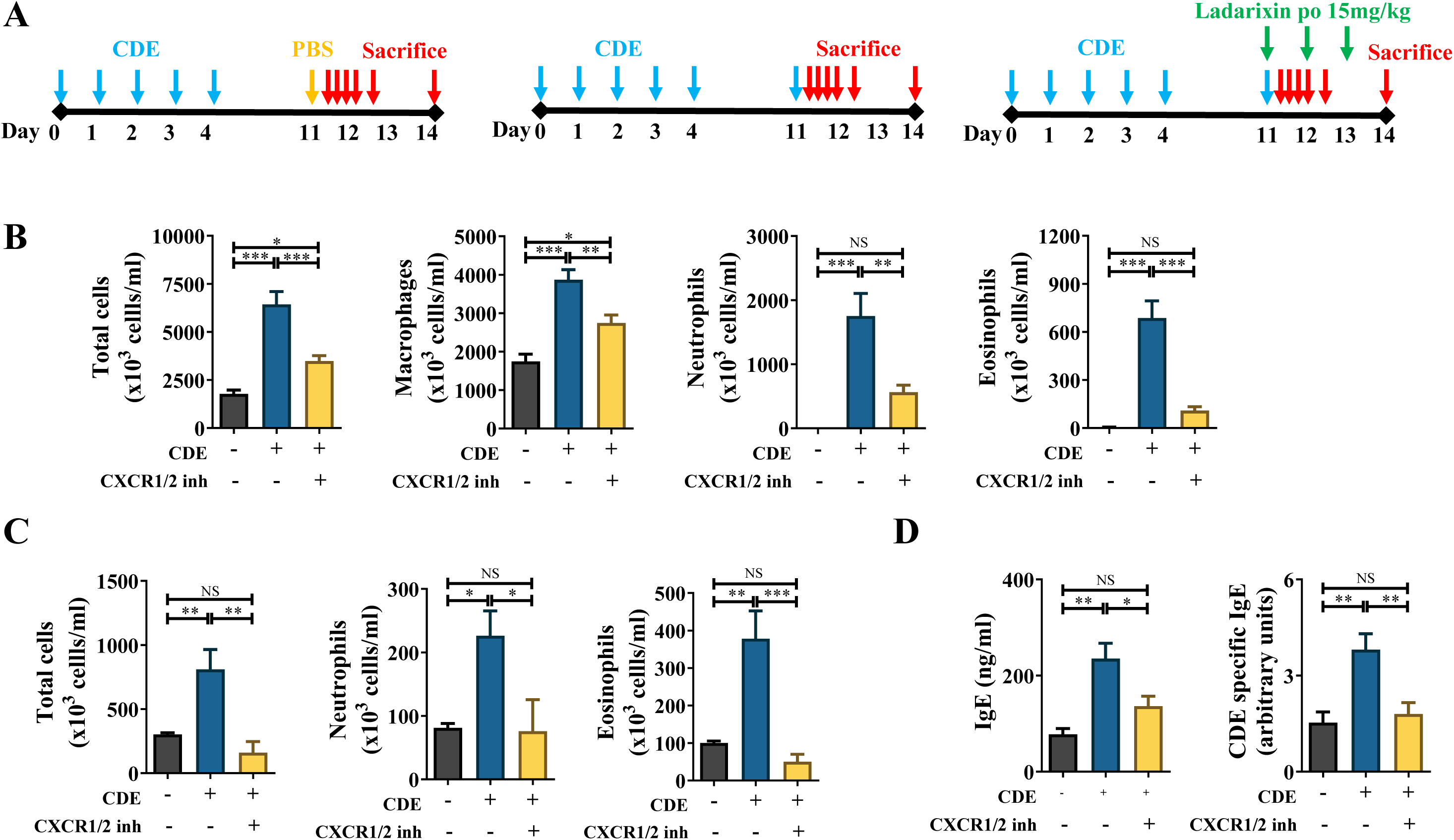
CXCR1 and CXCR2 stimulate allergic lung inflammation. (**A**) Protocol for CDE-MCM with or without ladarixn treatment. (**B**) The numbers of inflammatory cells in BALF at 72 h post-CDE challenge in CDE-MCM. (**C**) FCM analysis of the numbers of inflammatory immune cells in the lungs at 72 h post-CDE challenge in CDE-MCM. (**D**) The levels of serum total IgE and CDE-specific IgE at 72 h post-CDE challenge in CDE-MCM. BALF; bronchoalveolar lavage fluid CDE; cat dander extract CDE-MCM; CDE-multiple challenge model CDE-SCM; CDE-single challenge model

### CDE Single Challenge Model (CDE-SCM) to induce innate lung inflammation without sensitization

WT mice were intranasally challenged with a single dose of 100 µg/60µl of CDE and administered orally 15 mg/kg body weight of ladarixin simultaneously and sacrificed 28 h post-CDE challenge.

### Ladarixin

GMP human-use grade ladarixin was used in all studies performed in this manuscript and was a gift from Dompe pharmaceutical company (Dompé farmaceutici, L’Aquila, Italy).

### Processing of mouse BALF and lung tissue samples

The BALF and lung were obtained as described previously^28^.

### Bone marrow cells isolation

Bone marrow cells were isolated from the femur and tibia bones of the mouse. After red blood cells lysis by red blood cell lysing buffer (Sigma-Aldrich, St. Louis, MO), the bone marrow cells were frozen for subsequent experiments.

### qRT-PCR analysis

Total RNA from mouse lungs, bone marrow cells, or BALF cells were extracted with an RNeasy kit (Qiagen, Valencia, CA). cDNA was synthesized using a cDNA Synthesis kit (Qiagen). Amplification by real-time PCR was performed on a CFX Connect Real-Time PCR Detection System (Bio-Rad Laboratories, Hercules, CA) using SYBR Green PCR Master Mix Kit (Bio-Rad Laboratories) to examine lung mRNA expression of *Cxcl1, Cxcl 2, Cxcl 3, Cxcl 4, Cxcl 5*, *Cxcl* 7, *Cxcr1*, *Cxcr2*. *Gata3, Il1b, Il13, Il17, Il23, Il4, Il5, Il6, Rorc*, and *Tgfb1.* These primers were purchased from Integrated DNA Technologies (Coralville, IA).

### Measurement of IL23 protein in the lung tissue

Mouse lungs were lysed with Cell Lysis Buffer (Cell Signaling Technology, Danvers, MA) and sonicated, then the content of IL23 in the lung lysate was measured using a DuoSet ELISA development kit (R&D Systems, Minneapolis, MN) according to the manufacturer’s protocol.

### Lung single cells isolation

Lung tissues from mice were cut into small pieces and incubated with liberase-thermolysin medium (Sigma-Aldrich) in 42 µg/mL and 20% heat-inactivated fetal bovine serum in Hanks balanced salt solution at 37°C for 20 mins. After passing the digested lung through a 70 µm cell strainer, ice-cold PBS was added to neutralize the enzyme activity. After several wash steps with PBS, the lung single cells were used for flow cytometry analysis or culture system.

### Flow cytometry analysis with lung single cells

Lung single cells were pre-incubated with TruStain FcX solution (BioLegend, San Diego, CA) for 10 mins at 4°C, and stained with fluorophore-conjugated antibodies Fixable Viability Dye eFluor 780 (eBioscience, San Diego, CA), CD45 monoclonal antibody-pacific orange (Thermo Fisher Scientific, MA, USA, #MCD4530, clone 30-F11), anti-mouse CD3-Alexa Fluor 700 (Biolegend, #100216, clone 17A2), rat anti-mouse CD4-Brilliant Violet 520 (BD Biosciences, #563106, clone RM4.5), PE rat anti-mouse CD181 (CXCR1) (BD Biosciences, San Jose, CA, #566383, clone U45-632), BV711 Rat Anti-Mouse CD182 (CXCR2), (BD Biosciences, #747812, clone V48-2310), BV786 rat anti-mouse Siglec-F (BD Biosciences, #740956, clone E50-2440), Spark YG 593 anti- mouse Ly-6G Antibody (Biolegend, #127668, clone 1A8), Spark NIR™ 685 anti-mouse/human CD11b Antibody (Biolegend, #101278, clone M1/17) at 1:100 dilution in flow stain buffer containing FBS for 30 mins at 4°C, then cells were permeabilized with Fixation/Permeabilization kit (BD Biosciences) according to the protocol from vendor and stained with intracellular cytokine specific antibodies Rat anti-mouse Rorγt- Brilliant Violet-650 (BD Biosciences, #564722, clone Q31-378), mouse anti-GATA3 Alexa Fluor F488 (BD Bioscience #560163, clone L50-823,) Rat anti-mouse IL4-PE/Cyanine7 (BD Biosciences, #560699, clone 11B11), anti-mouse/anti-human IL5-Allophycocyanin (APC) (BD Biosciences, #554396, clone TRFK5), IL13 monoclonal antibody ((eBio13A), Brilliant Ultra Violet 805, eBioscience), Rat Anti-Mouse IL17A-PE (BD Biosciences, # 559502, clone TC11-18H10), and rat anti-mouse IL23 p19 Alexa Fluor 647 (BD Biosciences, # 565317, clone N71-1183) for 45 mins at 4°C. After washing, a flow cytometer was performed using a high-parameter Cytek®Aurora Flow Cytometer. Staining specificity was determined by fluorescence minus one (FMO) control to enhance the reliability of the gating analysis. Absolute cell numbers were quantified using Precision true count beads (BioLegend). The flow cytometry data was analyzed using FlowJo 10.8.1 Software. FC plots showing gating strategies and FMO are shown in **Supplemental Figure 3**. Neutrophils were identified as live CD45+ CD11b+ Ly6G+ Siglec F- cells. Eosinophils were identified as live CD45+ CD11b+ Siglec F+ Ly6G- cells. CD4+ T-cells were identified as live CD45+ CD3+ CD4+ cells.

### Measurement of serum total IgE and CDE-specific IgE

The methods have been described previously ^28^. Briefly, the plates were coated with CDE overnight or rat anti-mouse IgE (BD Biosciences) for 2 h. After blocking with sea block buffer for 2 h, the serum from the mice were added onto the plate. After washing, biotin-conjugated rat anti-mouse IgE (BD Biosciences) were plated onto the plate and incubated with avidin-conjugated alkaline phosphatase for 45 mins at 4°C (Sigma-Aldrich). Fluorescence intensities were measured with AttoPhos Substrate Solution (Promega, Madison, WI) using the Varioskan LUX reader (Thermo Fisher Scientific).

### Culture of lung single cells with chemokine cocktail

Lung single cells from sensitized mice were incubated with or without chemokine cocktail (CXCL1, CXCL2, CXCL3, and CXCL6 at 250 ng/ml each) for 3 days at 37 °C in the presence or absence of 1 h-pretreatment with ladarixin at 10uM. On day 4, after washing the cells, the cells were pre-incubated with Brefeldin A for 1 h, then, treated in the same way as above. On day 5, the cells were applied for flow cytometry.

### Statistical Analysis

The statistical analysis was performed by unpaired t-test for comparison of two groups or ANOVA for three or more groups using the software package GraphPad Prism 6 (GraphPad Software, San Diego, CA). The results are shown as mean ± SEM. All statistical analyses indicated data as significant at p < 0.05. *=P< .05, **=P< .01, ***=P< .001, ****=P< .0001.

## Results

### CDE challenge increases CXCL mRNA expression in the lung cells and CXCR1/2 in bone marrow cells of sensitized mice

Mice were sensitized to CDE by subjecting them to CDE-MCM (**Figure 1A**) ^26,28^. Compared to PBS challenge (**Figure 1A** left panel), CDE challenge (**Figure 1A** middle panel) in these sensitized mice induced much greater lung expression levels of *Cxcl1*, *Cxcl2*, *Cxcl3*, and *Cxcl5* at 2 h and/or 4 h, but not *Cxcl4* and *Cxcl7* (**Supplemental Figure 1A**). A shared feature of these upregulated *Cxcl1*, *Cxcl2*, *Cxcl3*, and *Cxcl5* chemokines is that they bind CXCR1 and CXCR2 ^29,30^. We reasoned that expression and secretion of these *Cxcl* chemokines in the lungs should increase in the number of progenitor cells expressing CXCR1 and CXCR2 ^29,30^ in the bone marrow to prepare them for migration to the lungs. To test this hypothesis, we examined the time kinetics of CDE challenge-induced upregulation of *Cxcr1* and *Cxcr2* mRNA expression in bone marrow of sensitized mice. CDE challenge increased the mRNA expression of *Cxcr1* and *Cxcr2* in bone marrow cells at 16 h post-CDE challenge (**Supplemental Figure 1B**), suggesting that these receptors play a role in recruiting inflammatory cells from bone marrow to the lungs.

### Allosteric inhibition of CXCR1 and CXCR2 blocks allergen challenge-induced allergic lung inflammation

Compared to PBS challenge, CDE challenge in CDE-MCM (**Figure 1A**) increased the number of total cells, macrophages, neutrophils, and eosinophils in BALF (**Figure 1B**) and lung tissues quantified by flow cytometry at 72 h after the final CDE challenge (**Figure 1C**), as well as total IgE and CDE-specific IgE in serum (**Figure 1D**). In the present study, allosteric inhibition of CXCR1 and CXCR2 was performed by oral administration of ladarixin, a small molecule that binds to the allosteric pocket of the trans-membrane region of both receptors with a 100-fold higher affinity than first-generation CXCR1 and CXCR2 inhibitors^31^. Allosteric inhibition of CXCR1 and CXCR2 in CDE-MCM model (**Figure 1A**, **right panel**) inhibited recruitment of macrophage into the BAL compartment (**Figure 1B**), and abrogated recruitment of neutrophils and eosinophils into BALF (**Figure 1B**) and lung compartments (**Figure 1C**).

### Allosteric inhibition of CXCR1 and CXCR2 blocks allergen challenge-induced recruitment of Th2-cells

The ability of allosteric inhibition of CXCR1 and CXCR2 to abrogate neutrophil and eosinophil recruitment made us ask the question that this inhibition may also affect Th2 and Th17-cells and genes. GATA3 is a key transcription factor associated with Th2-mediated allergic lung inflammation^3,4^. We examined *Gata3* mRNA expressions in BALF cells after CDE challenge in CDE-SCM and CDE-MCM (**Figure 1A**) models. Compared to naïve mice, CDE- challenge in CDE-SCM failed to increase mRNA expression levels of *Gata3* in BALF cells. By contrast, CDE challenge in CDE-MCM sensitized mice induced higher levels of *Gata3* mRNA expression in BALF cells compared to naïve mice and the mice subjected to CDE-SCM (**Figure 2A**). Allosteric inhibition of CXCR1 and CXCR2 in mice subjected to CDE-MCM (**Figure 1A right panel**) abrogated CDE challenge-induced increase in *Gata3* mRNA expression in BALF cells (**Figure 2B**). CDE challenge in CDE-MCM model upregulated expression levels of *Il4, Il5*, *Il13*, and *Tgfb1* in BALF cells (**Figure 2C**), expression levels of *Il4* and *Il13* in lung tissue (**Figure 2D**), and recruited IL4, IL5, IL13-secreting and GATA3-expressing Th2-cells into the lungs (**Figure 2E**). Allosteric inhibition of CXCR1 and CXCR2 abrogated each of these effects of CDE challenge (**Figure 2C-E**).

**Figure 2.**
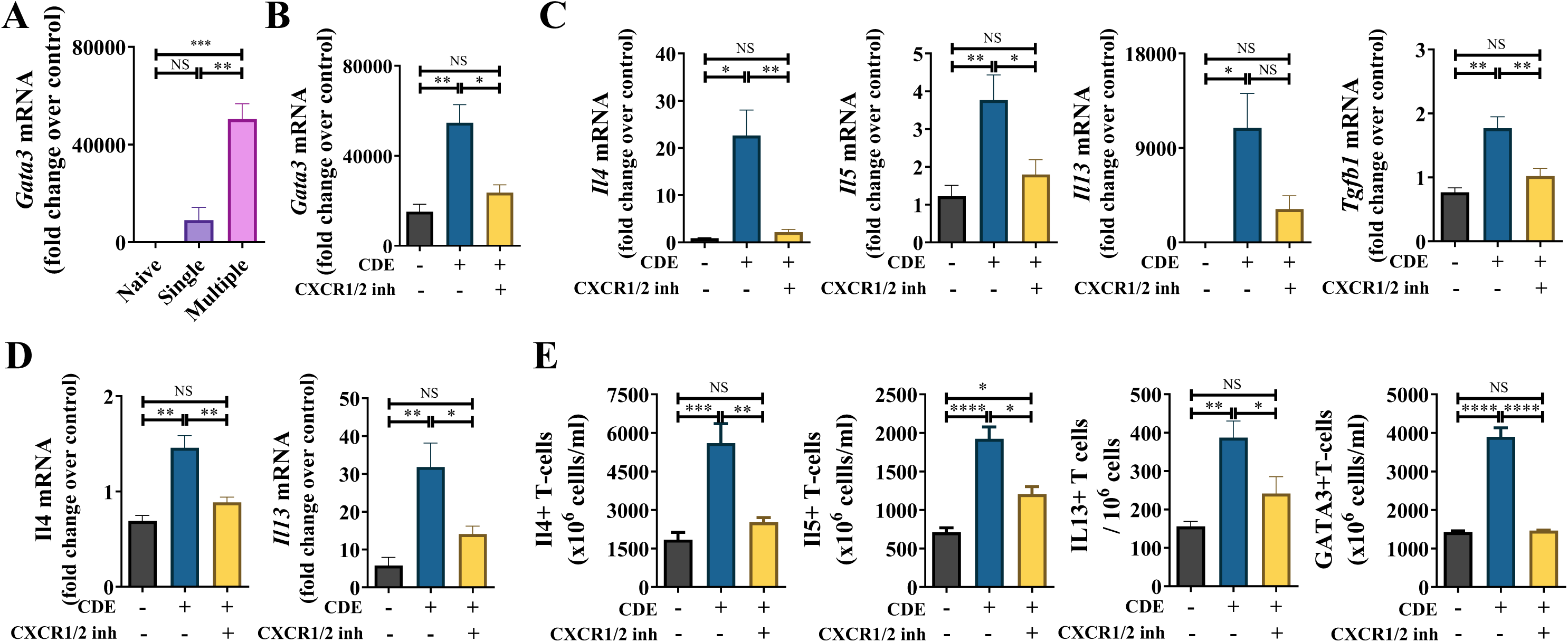
CXCR1 and CXCR2 recruit Th2-cells. (**A**) *Gata3* mRNA expressions in the BALF cells after CDE challenge in CDE-SCM and CDE-MCM mice at 28 h post-CDE challenge. (**B-E**) at 28 h post-CDE challenge in CDE-MCM. (**B**) *Gata3* mRNA expressions in BALF cells. (**C**) *Il4, Il5, Il13*, and *Tgfb1* cells mRNA expressions in BALF cells at 28h after CDE challenge (**D**) The lung mRNA expressions of *Il4* and *Il13* at 28h post-CDE challenge. (**E**) The number of IL4, IL5, IL13, and GATA3 expressing lung Th2-cells at 72h post-CDE challenge. BALF; bronchoalveolar lavage fluid CDE; cat dander extract CDE-MCM; CDE-multiple challenge model GATA3; GATA binding protein 3 Th2; T helper 2

### Allosteric inhibition of CXCR1 and CXCR2 blocks allergen challenge-induced Th17- associated mRNA expression and recruitment of Th17-cells

IL23 is a crucial cytokine that regulates the survival of Th17-cells and mediates Th17 inflammation^32^. Prior studies have shown that IL23, IL1β, and IL6 are elevated in the BALF obtained from the subjects with asthma ^33,34^ and these cytokines stimulate Th17 differentiation ^5–7^ by upregulating its master regulator RORγt^8^. Building on our observation that Allosteric inhibition of CXCR1 and CXCR2 abrogates Th2-associated mRNA expression levels and recruitment of Th2-cells, we examined the effects of CDE challenge in sensitized mice (**Figure 1A**) on the expression levels of IL23, IL1β, and IL6. CDE challenge upregulated *Il23, Il1b,* and *Il6* mRNA expression in BALF cells (**Figure 3A**) and lung tissues (**Figure 3B**). CDE challenge increased IL23 protein level in lung tissues (**Figure 3C**), and the numbers of IL23 positive T- cells, neutrophils, eosinophils, and lung epithelial cells in single cells obtained from lung tissues (**Figure 3D**). Allosteric inhibition of CXCR1 and CXCR2 abrogated each of these effects of CDE challenge (**Figure 3A-D**).

**Figure 3.**
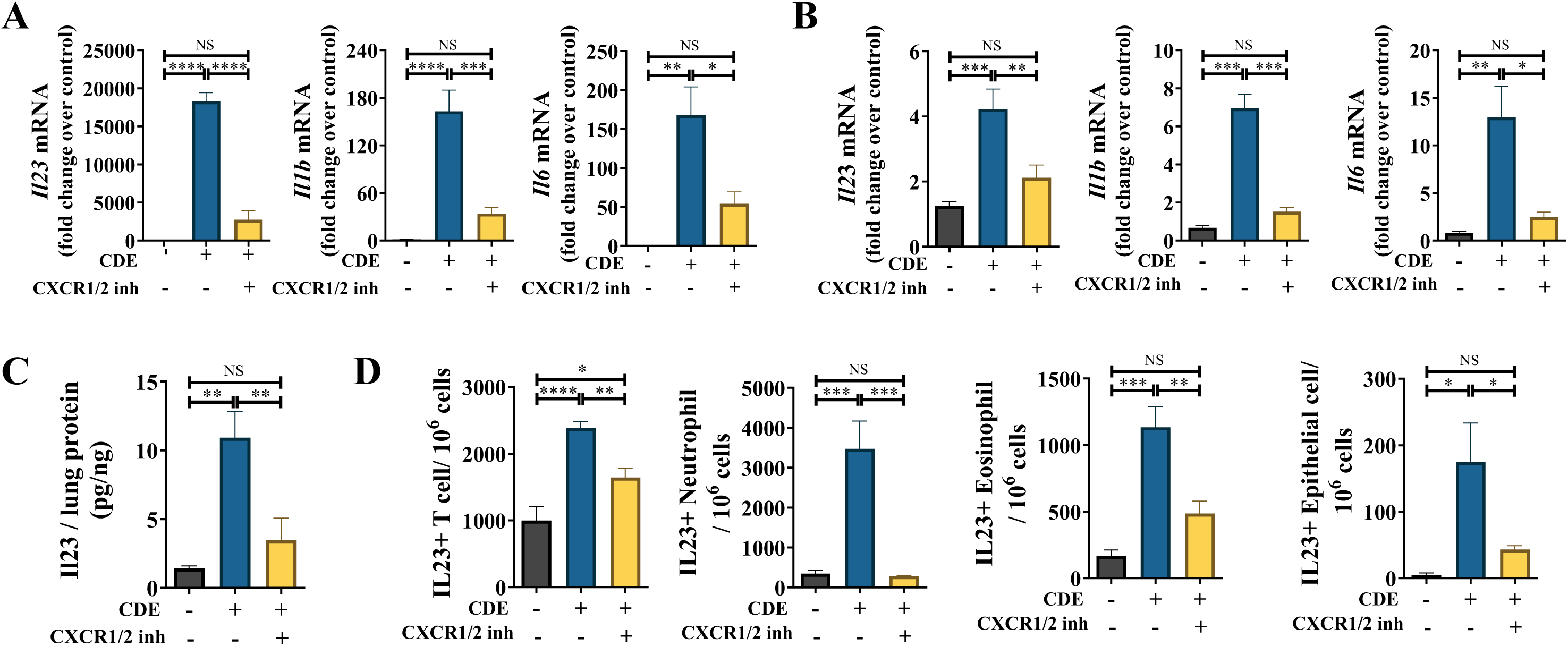
CXCR1 and CXCR2 upregulate Th17-associated mRNA and protein expression. (**A-B**) *Il23, Il1b,* and *Il6* mRNA expressions in the BALF cells (**A**) and in the lung (**B**) in CDE-MCM at 28 h post-CDE challenge. (**C**) The protein levels of Il23 in the lung in CDE-MCM at 28 h post-CDE challenge. (**D**) The numbers of IL23 producing T-cells, neutrophils, eosinophils, and epithelial cells in the lung in CDE-MCM at 72 h post-CDE challenge. BALF; bronchoalveolar lavage fluid CDE; cat dander extract CDE-MCM; CDE-multiple challenge model Th17; T helper 17

Because inhibiting CXCR1 and CXCR2 reduced Th17-promoting cytokines, we next evaluated the ability of CDE challenge in recruiting Th17-cells to the lungs. Compared to naïve mice, CDE-challenge in CDE-SCM failed to increase expression levels of *Rorc* mRNA in BALF cells (**Figure 4A**). By contrast, CDE challenge in CDE-MCM sensitized mice induced higher levels of *Rorc* mRNA expression mRNA in BALF cells compared to naïve mice and mice subjected to CDE-SCM (**Figure 4A**). CDE challenge in CDE-MCM upregulated *Il17* and *Rorc* mRNA expression in BALF cells (**Figure 4B**), upregulated *Il17* mRNA in lung tissues (**Figure 4C**), increased the number of RORγt-expressing T-cells (**Figure 4D**), and IL17-expressing Th17-cells (**Figure 4E**). Allosteric inhibition of CXCR1 and CXCR2 abrogated upregulation of *Il17* and *Rorc* mRNA expression in BALF cells, and inhibited *Il17* mRNA expression in lungs and recruitment of RORγt and IL17- expressing CD4+ T-cells (**Figure 4B-E**).

**Figure 4.**
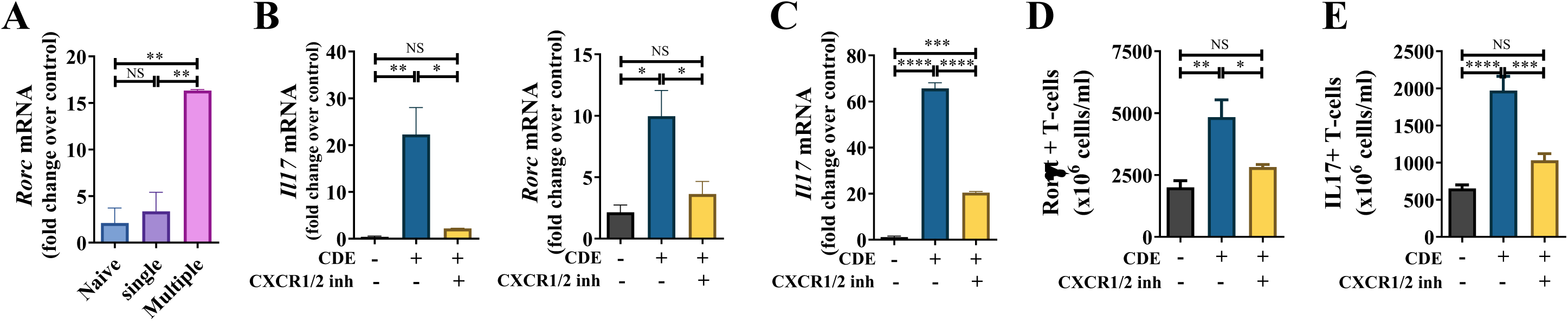
CXCR1 and CXCR2 recruit Th17-cells. (**A**) *Rorc* mRNA expressions in the BALF cells after CDE challenge in CDE-SCM and CDE-MCM mice at 28 h post-CDE challenge. (**B**) *Il17* and *Rorc* mRNA expressions in the BALF cells in CDE-MCM at 28 h post-CDE challenge. (**C**) *Il17* mRNA expressions in the lung in CDE-MCM at 28 h post-CDE challenge. (**D-E**) The number of RorγT T-cells (**D**) and IL17 secreting Th17-cells (**E**) in the lung in CDE-MCM at 72 h post-CDE challenge. BALF; bronchoalveolar lavage fluid CDE; cat dander extract CDE-MCM; CDE-multiple challenge model RorγT; retinoic acid–related orphan receptor-gamma t Th17; T helper 17

### Inhibition of CXCR1 and CXCR2 attenuates the recruitment of allergic inflammatory cells to the lungs

Neutrophils express CXCR1 and CXCR2 ^35^. Based on our observations showing the remarkable efficacy of allosteric inhibition of CXCR1 and CXCR2 in blocking CDE-challenge-induced recruitment of inflammatory cells (**Figure 2-4**), we hypothesized that various inflammatory cells recruited by CDE challenge also express CXCR1 and CXCR2. To test this hypothesis, we quantified the number of CXCR1 and CXCR2 expressing inflammatory cells in the lung in CDE-MCM that produce cytokines and key Th2 and Th17 transcription factors. CDE challenge in mice subjected to CDE-MCM increased the recruitment of CXCR1-expressing (**Figure 5A**) and CXCR2-expressing (**Figure 5B**) GATA3-, IL4-, IL5-, IL13-, RORγt-, and IL17- expressing T-cells. Furthermore, CDE challenge increased the recruitment of CXCR1- expressing (**Figure 5A**) and CXCR2-expressing (**Figure 5B**) neutrophils, eosinophils, IL23- expressing neutrophils, and IL23-expressing eosinophils. Allosteric inhibition of CXCR1 and CXCR2 abrogated CDE challenge-induced recruitment of CXCR1-expressing (**Figure 5A**) GATA3-, IL4-, IL5-, IL13-, RORγt-, and IL17-expressing T-cells, neutrophils, IL23-expressing neutrophils, and IL23-expressing eosinophils, and attenuated recruitment of CXCR1- expressing eosinophils. Likewise, allosteric inhibition of CXCR1 and CXCR2 abrogated recruitment of CXCR2-expressing IL4 and IL13 expressing T-cells, CXCR2-expressing neutrophils, IL23-expressing neutrophils, eosinophils, IL23-expressing eosinophils (**Figure 5B**), and attenuated recruitment of GATA-expressing T-cells, RORγt expressing and IL17- expressing T-cells. Together these results indicate that CXCR1 and CXCR2 are key regulators of Th2 and Th17-mediated allergic inflammation.

**Figure 5.**
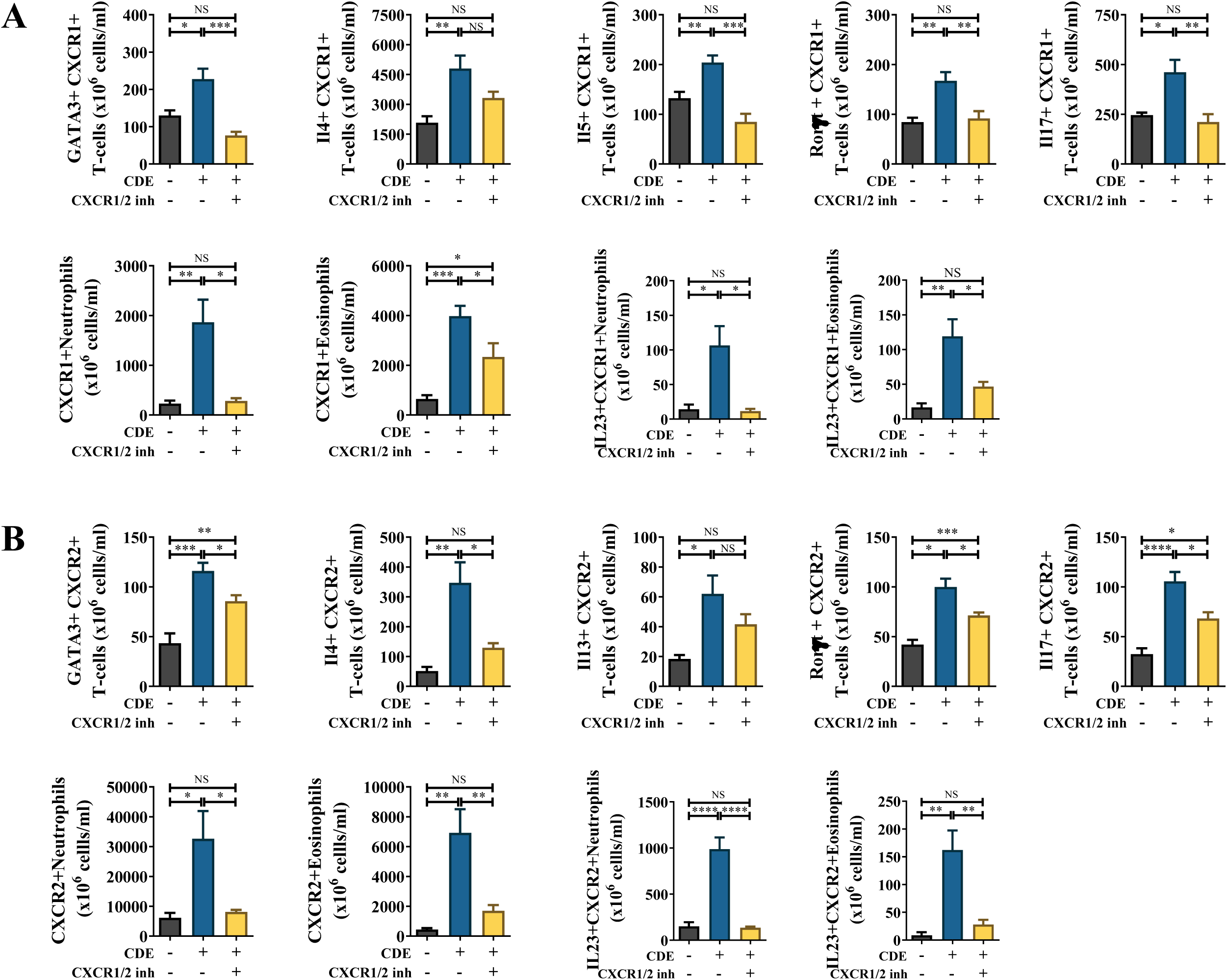
CXCR1 and CXCR2 recruit allergic inflammatory cells. (**A-B**) CDE-MCM at 72 h post-CDE challenge. (**A**) The numbers of CXCR1 expressing cells in the lung. (**B**) The numbers of CXCR2 expressing cells in the lung. CDE; cat dander extract CDE-MCM; CDE-multiple challenge model

### CXCR1 and CXCR2 stimulates proliferation of lung Th2- and Th17-cells

Th2-cells and Th17-cells regulate inflammation by releasing Th2 cytokines and Th17 cytokines followed by the recruitment of eosinophils ^3,4^ and neutrophils ^9^ in allergic airway inflammation. Building on our observations that CDE challenge in CDE-MCM **(Figure 1A**) upregulates *Cxcl1, Cxcl2, Cxcl3* and *Cxcl5* (**Supplemental Figure 2A**), we hypothesized that CXCL-CXCR1 and CXCR2 pathways may stimulate Th2-cells and Th17-cells. Lung single cells isolated from CDE-MCM mice were stimulated with a cocktail of CXCL1, CXCL2, CXCL3, and CXCL5 chemokines in the presence or absence of ladarixin. The CXCL chemokines increased the number of T cells (**Supplemental Figure 3A**), and IL4, IL5, and IL13-expressing Th2-cells and IL17-expressing Th17-cells (**Supplemental Figure 3B**). Inhibiting CXCR1 and CXCR2 by allosteric inhibition of CXCR1 and CXCR2 abrogated to cultured cells vigorously blocked this proliferation (**Supplemental Figure 3A-2B**). These data suggest that CXCR1 and CXCR2 drive CXCL-mediated proliferation of Th2 and Th17-cells.

## Discussion

Our findings indicate that CXCR1 and CXCR2 play a major role in Th2 and Th17- mediated allergic lung inflammation and proliferation. These receptors play a critical role in the recruitment of CXCR1- and CXCR2-expressing Th2-cells, Th17-cells, eosinophils, and neutrophils. We show for the first time that allosteric inhibition of CXCR1 and CXCR2 potently inhibits Th2 and Th17-mediated allergic lung inflammation. A growing number of monoclonal antibodies that target either a single Th2 cytokine or its cognate receptor are currently used to treat patients with severe asthma. Thus monoclonal antibodies block IL5 (mepolizumab^36^ and reslizumab^37^), IL5 receptor alpha-chain (benralizumab^38,39^), and IL4 receptor alpha (dupilumab^40^) are currently used in the management of severe asthma. IL17-secreting Th17- cells also contribute to airway remodeling and disease severity in severe endotype asthma^2^.

Biologics that block IL17 have been or are being tried in this asthma phenotype^41^. Unexpectedly, administration of the anti-IL17 receptor antibody brodalumab to the subjects with moderate to severe asthma failed to improve the clinical symptoms^41^. A number of studies indicate a crucial role of IL23 in maintaining Th17-cells ^32^, and regulating Th2- and Th17- cells^42,43^. Thus lack of IL23 suppressed naïve Th cell differentiation toward Th2-cells^42^, and treatment with anti-IL23 antibody suppressed ovalbumin-induced IL5 and IL13 production and eosinophil recruitment^43^. In the Th2/Th17 dominant subgroup of severe asthma^33,34^, BALF levels of IL23 were elevated^34^. These studies suggest that inhibiting IL23 with a monoclonal antibody should be beneficial in asthma. However, a clinical trial of human anti–IL23p19 monoclonal antibody risankizumab demonstrated that it had no beneficial effect in patients with severe asthma^44^. Thus, at this time, there is no effective treatment for Th17-associated allergic lung inflammation. Our results suggest that allosteric inhibition of CXCR1 and CXCR2 may provide a unique solution to suppress Th17-associated allergic inflammation. Our observations build on earlier reports showing that CXCR2 signaling regulates proliferation of megakaryocyte^45^, T-cell^46^, human melanoma cell^47^, and human pluripotent stem cells^48^.

A focus of earlier studies of CXCR1/2 inhibitors in mitigating neutrophil recruitment in allergic inflammation^25,26^. In addition to our novel finding that allosteric inhibition of CXCR1 and CXCR2 blocks the recruitment of Th2- and Th17-cells, we show that ladarixin suppresses the recruitment of neutrophils, including IL23-secreting neutrophils. These data build on our earlier report that neutrophils can augment allergic airway inflammation and sensitization^25^. Other studies have shown that neutrophil recruitment to the site is closely associated with local allergic inflammation in atopic dermatitis^49^, allergic contact dermatitis^50^, anaphylaxis^51^, and asthma ^25,52^ by regulating innate and adaptive immunity. Since neutrophils express abundant CXCR1 and CXCR2, inhibition of their recruitment likely also contributes to ladarixin-induced mitigation of allergic lung inflammation.

Ladarixin is a dual CXCR1/2 antagonist that demonstrated beneficial effects on tumors and endocrine disorders. Treatment with ladarixin inhibited the melanoma xenografts *in vivo*^31^, delayed and prevented spontaneous diabetes onset in NOD mice^53^, and when administered with an immune checkpoint inhibitor like anti-PD-1 inhibited tumor growth and improved the survival in tumor xenograft mouse model^54^. Based on the clinical trial recently performed in patients with new-onset type 1 diabetes, the use of ladarixin is safe and well tolerated^55^. We suggest that future human studies should carefully investigate the role of allosteric inhibition of CXCR1 and CXCR2 are a novel approach to mitigating allergic inflammation involving activation of Th2-and Th17-cells and pathways.

## Acknowledgments

We thank the Cytometry and Cell Sorting Core (CCSC) at Baylor College of Medicine for expert assistance.

This research was supported in parts by Department of Defense (grant no. PR171425 W81XWH-18-1-0743) and the National Heart, Lung, and Blood Institute (grant nos. 5R01HL145477-02 and 3R01HL145477-01S1).

## Supplemental Figure

**Supplemental Figure 1. Allergen challenge upregulates CXCLs and CXCR1/CXCR2**

**(A)** Cxcl expressions in the lungs at 2 h and 4 h post-CDE challenge in CDE-MCM. (**B**) Time kinetics of Cxcr1 and Cxcr2 expressions in the bone marrow cells in CDE-MCM.

CDE; cat dander extract

CDE-MCM; CDE-multiple challenge model

**Supplemental Figure 2. CXCR1/2 stimulates proliferation of Th2 and Th17-cells**

**(A)** The number of T-cells in culture lung cells. (**B**) The numbers of IL4, IL5, IL13, and IL17, and producing T-cells in culture lung cells.

**Supplemental Figure 3. Representative FC plot gating strategy**

The forward and side scatter (FSC vs SSC) density plots were generated to separate cell populations of interest from debris. Next, doublets and debris were excluded by forward scatter height versus forward scatter area density plot (FSC-H vs FSC-A). Next, dead cells were excluded by gating only live cells using fluorophore-conjugated antibodies Fixable Viability Dye eFluor 780. In all subsequent steps, side scatter area (SSC-A) was plotted against fluorescence- tagged antibodies to gate positive cells by excluding cells in the corresponding FMO gate. Sequentially, live cells were gated using CD45+ cells to gate hematopoietic cells, CD3+, CD4+ double positive cells to gate for CD4+ T-cells, CXCR1+ cells to gate CXCR1-expressing CD4+ T-cells, then intracellular IL4+ to gate for CXCR1+ IL4+ Th2-cells in the lungs.

FMO; fluorescence minus one

## References

1. Woodruff PG, Modrek B, Choy DF, et al. T-helper type 2-driven inflammation defines major subphenotypes of asthma. Am J Respir Crit Care Med. 2009;180(5):388–395.

2. Al-Ramli W, Prefontaine D, Chouiali F, et al. T(H)17-associated cytokines (IL-17A and IL-17F) in severe asthma. J Allergy Clin Immunol. 2009;123(5):1185–1187.

3. Zheng W, Flavell RA. The transcription factor GATA-3 is necessary and sufficient for Th2 cytokine gene expression in CD4 T cells. Cell. 1997;89(4):587–596.

4. Zhang DH, Yang L, Cohn L, et al. Inhibition of allergic inflammation in a murine model of asthma by expression of a dominant-negative mutant of GATA-3. Immunity. 1999;11(4):473–482.

5. Mangan PR, Harrington LE, O’Quinn DB, et al. Transforming growth factor-beta induces development of the T(H)17 lineage. Nature. 2006;441(7090):231-234.

6. Veldhoen M, Hocking RJ, Atkins CJ, Locksley RM, Stockinger B. TGFbeta in the context of an inflammatory cytokine milieu supports de novo differentiation of IL-17- producing T cells. Immunity. 2006;24(2):179–189.

7. Manel N, Unutmaz D, Littman DR. The differentiation of human T(H)-17 cells requires transforming growth factor-beta and induction of the nuclear receptor RORgammat. Nat Immunol. 2008;9(6):641–649.

8. Ivanov, II, McKenzie BS, Zhou L, et al. The orphan nuclear receptor RORgammat directs the differentiation program of proinflammatory IL-17+ T helper cells. Cell. 2006;126(6):1121–1133.

9. Miyamoto M, Prause O, Sjostrand M, Laan M, Lotvall J, Linden A. Endogenous IL-17 as a mediator of neutrophil recruitment caused by endotoxin exposure in mouse airways. J Immunol. 2003;170(9):4665–4672.

10. Sokol CL, Luster AD. The chemokine system in innate immunity. Cold Spring Harb Perspect Biol. 2015;7(5).

11. Kampen GT, Stafford S, Adachi T, et al. Eotaxin induces degranulation and chemotaxis of eosinophils through the activation of ERK2 and p38 mitogen-activated protein kinases. Blood. 2000;95(6):1911–1917.

12. Bonecchi R, Bianchi G, Bordignon PP, et al. Differential expression of chemokine receptors and chemotactic responsiveness of type 1 T helper cells (Th1s) and Th2s. J Exp Med. 1998;187(1):129–134.

13. Anderson CA, Patel P, Viney JM, Phillips RM, Solari R, Pease JE. A degradatory fate for CCR4 suggests a primary role in Th2 inflammation. J Leukoc Biol. 2020;107(3):455–466.

14. Lin R, Choi YH, Zidar DA, Walker JKL. beta-Arrestin-2-Dependent Signaling Promotes CCR4-mediated Chemotaxis of Murine T-Helper Type 2 Cells. Am J Respir Cell Mol Biol. 2018;58(6):745–755.

15. Panina-Bordignon P, Papi A, Mariani M, et al. The C-C chemokine receptors CCR4 and CCR8 identify airway T cells of allergen-challenged atopic asthmatics. J Clin Invest. 2001;107(11):1357–1364.

16. Acosta-Rodriguez EV, Rivino L, Geginat J, et al. Surface phenotype and antigenic specificity of human interleukin 17-producing T helper memory cells. Nat Immunol. 2007;8(6):639–646.

17. Annunziato F, Cosmi L, Santarlasci V, et al. Phenotypic and functional features of human Th17 cells. J Exp Med. 2007;204(8):1849–1861.

18. Wang C, Kang SG, Lee J, Sun Z, Kim CH. The roles of CCR6 in migration of Th17 cells and regulation of effector T-cell balance in the gut. Mucosal Immunol. 2009;2(2):173–183.

19. Ordonez CL, Shaughnessy TE, Matthay MA, Fahy JV. Increased neutrophil numbers and IL-8 levels in airway secretions in acute severe asthma: Clinical and biologic significance. Am J Respir Crit Care Med. 2000;161(4 Pt 1):1185-1190.

20. Qiu Y, Zhu J, Bandi V, Guntupalli KK, Jeffery PK. Bronchial mucosal inflammation and upregulation of CXC chemoattractants and receptors in severe exacerbations of asthma. Thorax. 2007;62(6):475–482.

21. Hosoki K, Ying S, Corrigan C, et al. Analysis of a Panel of 48 Cytokines in BAL Fluids Specifically Identifies IL-8 Levels as the Only Cytokine that Distinguishes Controlled Asthma from Uncontrolled Asthma, and Correlates Inversely with FEV1. PLoS ONE. 2015;10(5):e0126035.

22. Lippert U, Zachmann K, Henz BM, Neumann C. Human T lymphocytes and mast cells differentially express and regulate extra- and intracellular CXCR1 and CXCR2. Exp Dermatol. 2004;13(8):520–525.

23. Thomas KM, Taylor L, Navarro J. The interleukin-8 receptor is encoded by a neutrophil- specific cDNA clone, F3R. J Biol Chem. 1991;266(23):14839–14841.

24. Farkas L, Hahn MC, Schmoczer M, et al. Expression of CXC chemokine receptors 1 and 2 in human bronchial epithelial cells. Chest. 2005;128(5):3724–3734.

25. Hosoki K, Aguilera-Aguirre L, Brasier AR, Kurosky A, Boldogh I, Sur S. Facilitation of Allergic Sensitization and Allergic Airway Inflammation by Pollen-Induced Innate Neutrophil Recruitment. Am J Respir Cell Mol Biol. 2016;54(1):81–90.

26. Hosoki K, Rajarathnam K, Sur S. Attenuation of murine allergic airway inflammation with a CXCR1/CXCR2 chemokine receptor inhibitor. Clin Exp Allergy. 2019;49(1):130–132.

27. Hosoki K, Aguilera-Aguirre L, Brasier AR, Kurosky A, Boldogh I, Sur S. Pollen-induced Innate Recruitment of Neutrophils Facilitates Induction of Allergic Sensitization and Airway Inflammation. Am J Respir Cell Mol Biol. 2015.

28. Hosoki K, Jaruga P, Itazawa T, et al. Excision release of 5?hydroxycytosine oxidatively induced DNA base lesions from the lung genome by cat dander extract challenge stimulates allergic airway inflammation. Clin Exp Allergy. 2018;48(12):1676–1687.

29. Palmqvist C, Wardlaw AJ, Bradding P. Chemokines and their receptors as potential targets for the treatment of asthma. Br J Pharmacol. 2007;151(6):725–736.

30. Borish LC, Steinke JW. 2. Cytokines and chemokines. J Allergy Clin Immunol. 2003;111(2 Suppl):S460-475.

31. Kemp DM, Pidich A, Larijani M, et al. Ladarixin, a dual CXCR1/2 inhibitor, attenuates experimental melanomas harboring different molecular defects by affecting malignant cells and tumor microenvironment. Oncotarget. 2017;8(9):14428–14442.

32. McGeachy MJ, Chen Y, Tato CM, et al. The interleukin 23 receptor is essential for the terminal differentiation of interleukin 17-producing effector T helper cells in vivo. Nat Immunol. 2009;10(3):314–324.

33. Irvin C, Zafar I, Good J, et al. Increased frequency of dual-positive TH2/TH17 cells in bronchoalveolar lavage fluid characterizes a population of patients with severe asthma. J Allergy Clin Immunol. 2014;134(5):1175–1186 e1177.

34. Liu W, Liu S, Verma M, et al. Mechanism of TH2/TH17-predominant and neutrophilic TH2/TH17-low subtypes of asthma. J Allergy Clin Immunol. 2017;139(5):1548–1558 e1544.

35. Cummings CJ, Martin TR, Frevert CW, et al. Expression and function of the chemokine receptors CXCR1 and CXCR2 in sepsis. J Immunol. 1999;162(4):2341–2346.

36. Ortega HG, Liu MC, Pavord ID, et al. Mepolizumab treatment in patients with severe eosinophilic asthma. N Engl J Med. 2014;371(13):1198–1207.

37. Castro M, Zangrilli J, Wechsler ME, et al. Reslizumab for inadequately controlled asthma with elevated blood eosinophil counts: results from two multicentre, parallel, double-blind, randomised, placebo-controlled, phase 3 trials. Lancet Respir Med. 2015;3(5):355–366.

38. FitzGerald JM, Bleecker ER, Nair P, et al. Benralizumab, an anti-interleukin-5 receptor alpha monoclonal antibody, as add-on treatment for patients with severe, uncontrolled, eosinophilic asthma (CALIMA): a randomised, double-blind, placebo-controlled phase 3 trial. Lancet. 2016;388(10056):2128–2141.

39. Bleecker ER, FitzGerald JM, Chanez P, et al. Efficacy and safety of benralizumab for patients with severe asthma uncontrolled with high-dosage inhaled corticosteroids and long-acting beta(2)-agonists (SIROCCO): a randomised, multicentre, placebo-controlled phase 3 trial. Lancet. 2016;388(10056):2115–2127.

40. Wenzel S, Ford L, Pearlman D, et al. Dupilumab in persistent asthma with elevated eosinophil levels. N Engl J Med. 2013;368(26):2455–2466.

41. Busse WW, Holgate S, Kerwin E, et al. Randomized, double-blind, placebo-controlled study of brodalumab, a human anti-IL-17 receptor monoclonal antibody, in moderate to severe asthma. Am J Respir Crit Care Med. 2013;188(11):1294–1302.

42. Peng J, Yang XO, Chang SH, Yang J, Dong C. IL-23 signaling enhances Th2 polarization and regulates allergic airway inflammation. Cell Res. 2010;20(1):62–71.

43. Wakashin H, Hirose K, Maezawa Y, et al. IL-23 and Th17 cells enhance Th2-cell-mediated eosinophilic airway inflammation in mice. Am J Respir Crit Care Med. 2008;178(10):1023–1032.

44. Brightling CE, Nair P, Cousins DJ, Louis R, Singh D. Risankizumab in Severe Asthma - A Phase 2a, Placebo-Controlled Trial. N Engl J Med. 2021;385(18):1669–1679.

45. Emadi S, Clay D, Desterke C, et al. IL-8 and its CXCR1 and CXCR2 receptors participate in the control of megakaryocytic proliferation, differentiation, and ploidy in myeloid metaplasia with myelofibrosis. Blood. 2005;105(2):464–473.

46. Lee YS, Won KJ, Park SW, et al. Mesenchymal stem cells regulate the proliferation of T cells via the growth-related oncogene/CXC chemokine receptor, CXCR2. Cell Immunol. 2012;279(1):1–11.

47. Shang FM, Li J. A small-molecule antagonist of CXCR1 and CXCR2 inhibits cell proliferation, migration and invasion in melanoma via PI3K/AKT pathway. Med Clin (Barc). 2019;152(11):425–430.

48. Jung JH, Lee SJ, Kim J, et al. CXCR2 and its related ligands play a novel role in supporting the pluripotency and proliferation of human pluripotent stem cells. Stem Cells Dev. 2015;24(8):948–961.

49. Oyoshi MK, He R, Li Y, et al. Leukotriene B4-driven neutrophil recruitment to the skin is essential for allergic skin inflammation. Immunity. 2012;37(4):747–758.

50. Weber FC, Nemeth T, Csepregi JZ, et al. Neutrophils are required for both the sensitization and elicitation phase of contact hypersensitivity. J Exp Med. 2015;212(1):15–22.

51. Jonsson F, Mancardi DA, Kita Y, et al. Mouse and human neutrophils induce anaphylaxis. The Journal of clinical investigation. 2011;121(4):1484–1496.

52. Hosoki K, Boldogh I, Aguilera-Aguirre L, et al. Myeloid differentiation protein 2 facilitates pollen- and cat dander-induced innate and allergic airway inflammation. J Allergy Clin Immunol. 2016;137(5):1506–1513 e1502.

53. Citro A, Valle A, Cantarelli E, et al. CXCR1/2 inhibition blocks and reverses type 1 diabetes in mice. Diabetes. 2015;64(4):1329–1340.

54. Piro G, Carbone C, Agostini A, et al. CXCR1/2 dual-inhibitor ladarixin reduces tumour burden and promotes immunotherapy response in pancreatic cancer. Br J Cancer. 2022.

55. Piemonti L, Keymeulen B, Gillard P, et al. Ladarixin, an inhibitor of the interleukin-8 receptors CXCR1 and CXCR2, in new-onset type 1 diabetes: A multicentre, randomized, double-blind, placebo-controlled trial. Diabetes Obes Metab. 2022.

