## Supplementary figures and images for "Allosteric inhibition of CXCR1 and CXCR2 abrogates Th2/Th17-associated Allergic Lung Inflammation in Mice"

### Supplemental Figure 1

#### Supplemental Figure 1

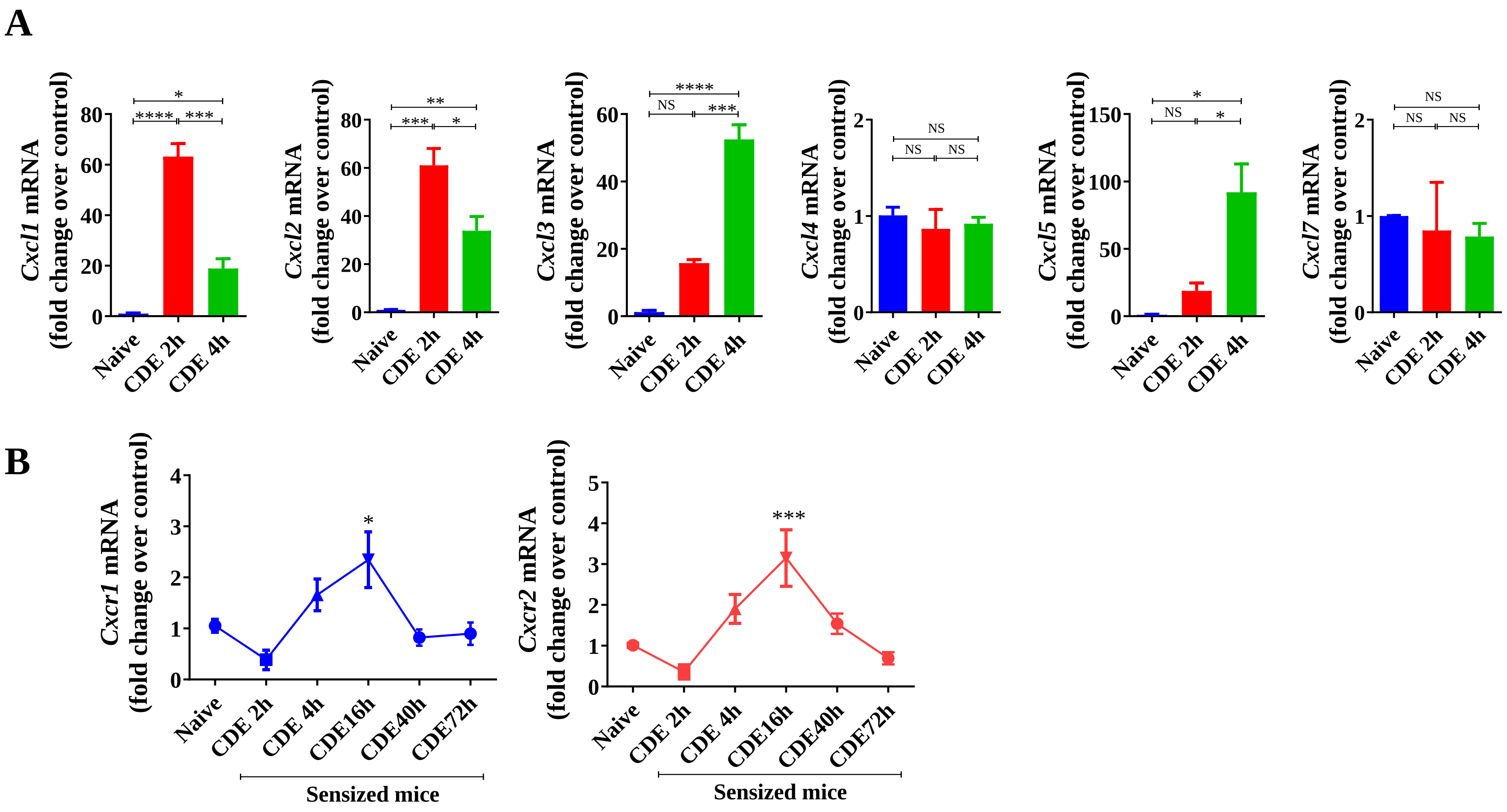

### Supplemental Figure 2

#### Supplemental Figure 2

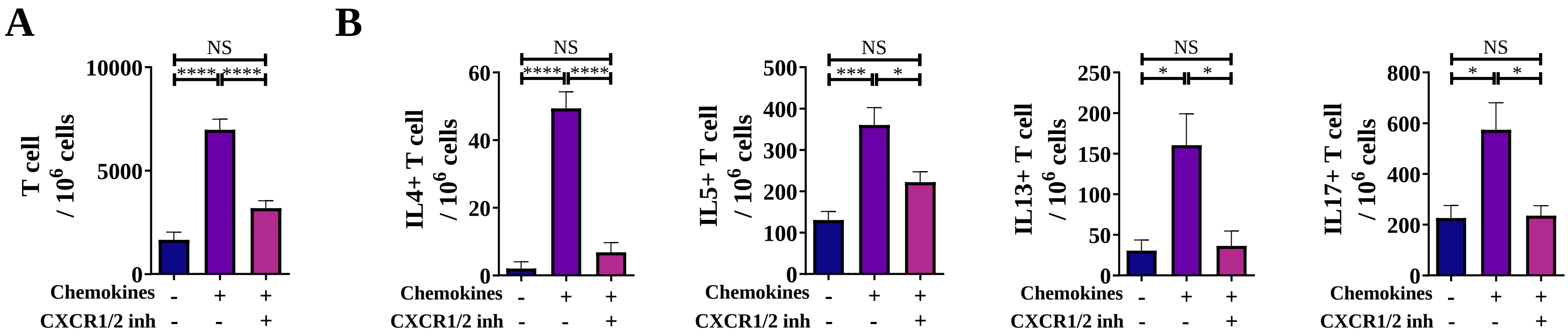

### Supplemental Figure 3

#### Supplemental Figure 3

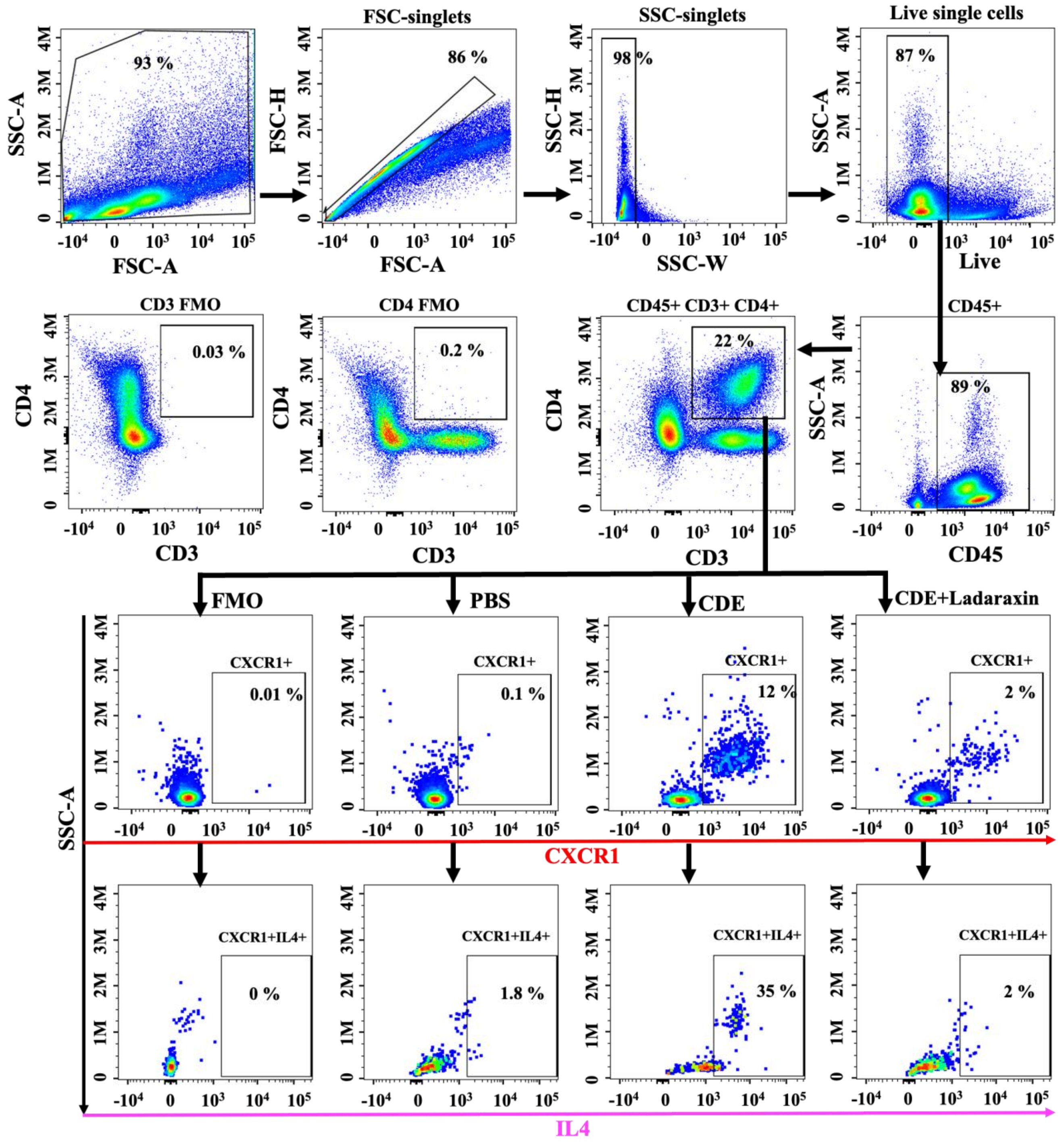
